# Paradoxical Tumor Suppressor Role of Yorkie Through a TOR‑Dependent α‑Tubulin Acetylation in Select Squamous Epithelium

**DOI:** 10.1101/2024.08.08.607173

**Authors:** Rachita Bhattacharya, Jeganath Ammavasai, Shruti Agarwal, Hamim Zafar, Pradip Sinha, Nitin Mohan

## Abstract

Protooncogenes, which can also function as tumor suppressors, are termed double agents. The Hippo pathway transcriptional coactivators YAP/TAZ and their *Drosophila* homolog Yorkie (Yki) are known oncogenes. Paradoxically, in squamous epithelia, they are constitutively active yet non-oncogenic and can even function as tumor suppressors. We previously found that loss of Yki in the *Drosophila* male accessory gland (MAG) squamous epithelium promotes squamous cell carcinoma (SCC) by upregulating TOR signaling, but the mechanism was unknown. Here, we show that this tumorigenesis stems from TOR-mediated hyperacetylation of α-tubulin. This pathway is tissue-specific, as the ovarian follicular epithelium is resistant to Yki loss. A comparison of MAG and follicular epithelia single-cell transcriptomic data revealed their distinct Yki-TOR signaling states, suggesting lineage-specific mechanoconstraints in these two lineages. Our results identify suppression of TOR-driven α-tubulin acetylation as the mechanistic basis for Yki’s role as a double-agent in select squamous epithelia.

## Introduction

Some protooncogenes can function as tumor suppressors in specific cell types. Examples include NOTCH1 (Lobry et al., 2014), TP53/p53 (Soussi and Wiman, 2015), or RB1(Chau and Wang, 2003). Such genes are described as “double agents” due to their context-specific opposing roles in carcinogenesis (Shen et al., 2018). YAP/TAZ in mammals, and Yorkie (Yki) in *Drosophila*, are also protooncogenes. Out-of-context activation of these proteins promotes cancer (Huang et al., 2005; Zanconato et al., 2016). However, in squamous epithelia, nuclear YAP/TAZ/Yki signaling is perpetual. This is due to cell flattening, which provides a mechanosensory relief from their cytoplasmic sequestration (Dupont et al., 2011; Fletcher et al., 2018; Kowalczyk et al., 2022; Nakajima et al., 2017). Such a scenario of developmentally programmed and chronic YAP/TAZ/Yki signaling likely occludes its protooncogenic roles in select squamous epithelia. For instance, loss of YAP/TAZ signaling results in lung squamous cell carcinoma (Gao et al., 2014; Huang et al., 2017), thereby revealing its tumor-suppressor-like role (Pearson et al., 2021)(for review, see (Baroja et al., 2024; Franklin et al., 2023; Luo et al., 2023)). Recently, we have shown that in *Drosophila*, too, loss of Yki signaling in the squamous epithelium of the male accessory gland (MAG) triggers TOR signaling and induces re-entry into the cell cycle, culminating in squamous cell carcinoma (SCC) (Bhattacharya et al., 2024). This outcome contrasts with the follicular squamous epithelium (Borreguero-Muñoz et al., 2019; Fletcher et al., 2018), where loss of Yki signaling fails to trigger SCC, even though nuclear Yki signaling is a common feature of both these epithelial types. Thus, Yki, like its human counterparts YAP/TAZ, displays the hallmarks of a double agent in select cell types. Yet the mechanistic basis of its paradoxical tumor suppressor activity—specifically how Yki restrains TOR-driven processes to prevent carcinoma in squamous epithelia—remains unresolved.

Several lines of recent evidence suggest a hitherto unexplored possibility of a tumor suppressor-like role of developmentally programmed nuclear Yki signaling in select squamous epithelia. For example, Hippo signaling is regulated by microtubule acetylation (Mok and Choi, 2022), which is required for cell stretching or flattening (Moise et al., 2024; Seetharaman et al., 2022) and also implicated in carcinogenesis (Wattanathamsan and Pongrakhananon, 2023). We therefore asked whether a Yki–TOR–microtubule acetylation axis explains MAG-SCC. Here, we test this hypothesis and uncover a lineage-specific mechanism in which suppression of TOR-driven α-tubulin acetylation underlies Yki’s paradoxical role as a double-agent tumor suppressor in *Drosophila* squamous epithelium.

## Results

### Microtubule acetylation is required for squamous cell flattening

Because epithelial stretching correlates with microtubule acetylation *in vitro*, we examined its endogenous levels during *Drosophila* follicular and MAG epithelial flattening. In both, the columnar-to-squamous transition coincided with elevated microtubule acetylation (**Fig. 1A–B and Fig. 1—figure supplement 1A-D**). In the ovarian follicular epithelium, for instance, total α-tubulin and acetylated microtubule levels rose in parallel during the columnar-to-squamous transition, suggesting near-complete acetylation of the tubulin pool in flattened squamous cells (**Fig. 1—figure supplement 2A–D**).

**Figure 1.**
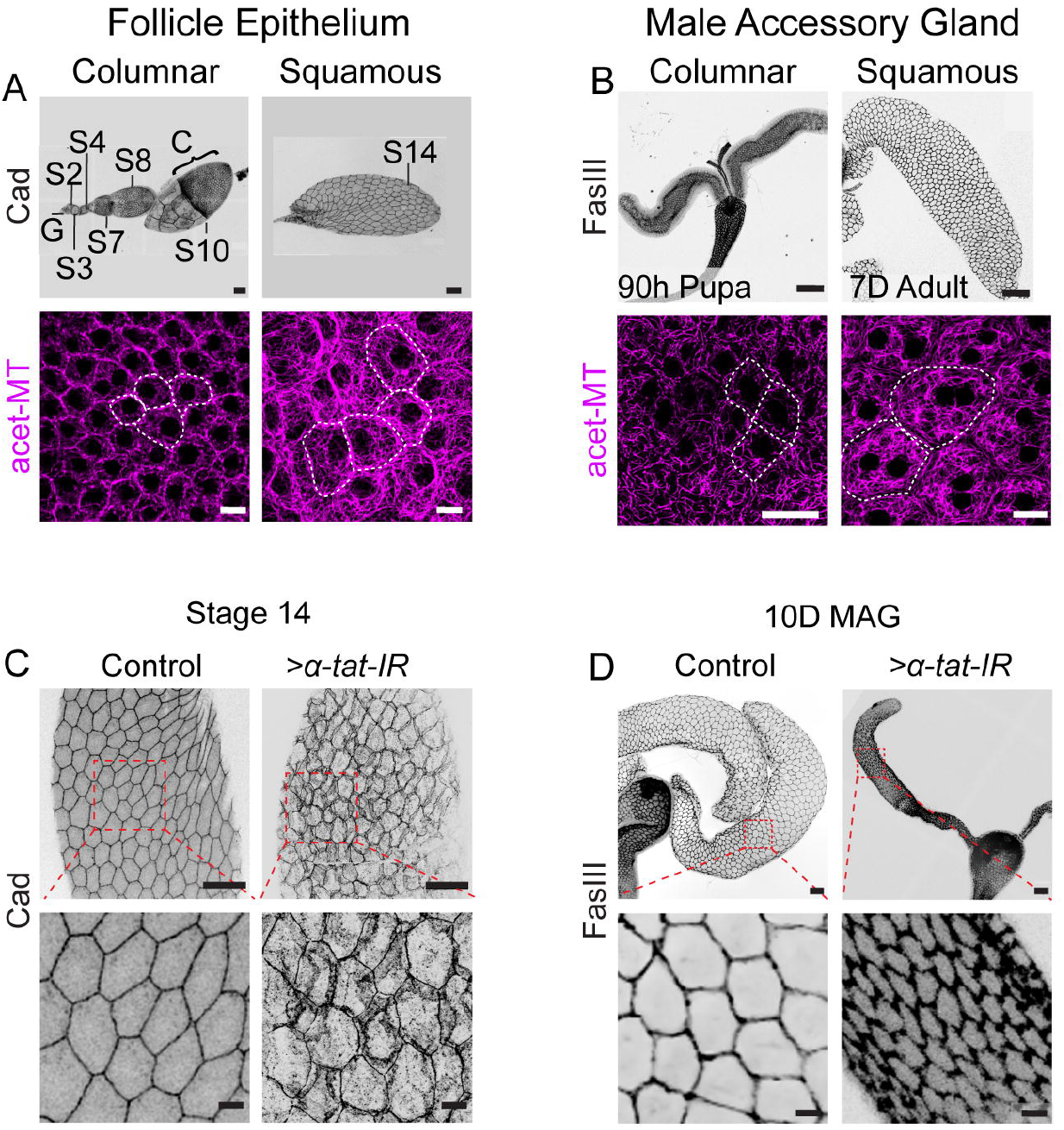
Microtubule acetylation is elevated in squamous epithelia and required for cell flattening. **(A)** Follicle epithelium development (Top). Ovariole showing the transition from columnar (Stage 10) to squamous (Stage 14) morphology, visualized with a Cadherin reporter (*shg-mTomato*). (Bottom) Immunostaining reveals increased acetylated α-tubulin levels in Stage 14 squamous cells compared to Stage 10 columnar cells. **(B)** Pupal (90h APF)-to-adult (7D) development of the male accessory gland (MAG) squamous epithelium, showing increased acetylated α-tubulin levels during columnar-to-squamous transition. (**C, D**) Knockdown of *α-tat* impairs cell flattening in both Stage 14 follicle cells (C, *GR1-Gal4>α-tat-IR*) and adult MAG epithelium (D, *ov-Gal4>α-tat-IR*). Insets magnified below. Abbreviation in this and all subsequent figures: G=germarium, S=Stage of oocytes, C=Columnar epithelium, MAG= male accessory gland, Acet-MT=acetylated microtubules, IR=RNAi, h=hours, APF=after pupa formation, D=days. Scale bars: upper panels (A–B) 100 µm; lower panels (A, B) and all insets 10 µm.

To test the functional significance of this acetylation, we knocked down *α-tat* (which encodes α-tubulin acetyltransferase). Loss of *α-tat* activity did not affect total α-tubulin organization or density (**Fig. 1—figure supplement 3A–B**), but reduced cell area in both follicular (*GR1-Gal4>α-tat-IR*, **Fig. 1C and Fig. 1—figure supplement 1E**) and MAG (*ov-Gal4>α-tat-IR*, **Fig. 1D and Fig. 1—figure supplement 1F**) squamous epithelia. Loss of cell flattening was accompanied by corresponding decreases in organ size in both the MAG and egg chambers (**Fig. 1D**, top panel; **Fig. 1—figure supplement 3C–D**). These results establish a foundation to explore how tubulin acetylation impacts additional cellular processes integral to squamous morphogenesis. Thus, we next investigated how microtubule acetylation affects membrane protein trafficking during epithelial flattening.

### α-tat loss disrupts junctional protein localization and actin organization

Cell flattening during squamous cell morphogenesis reduces cell–cell adhesion (Gomez et al., 2012). We also noticed low levels of the lateral membrane protein FasIII (Wells et al., 2013), the sub-apical adherens junction protein, DE-Cadherin (Huang et al., 2012; Pope and Harris, 2008), and the basolateral protein, Lgl (Bilder et al., 2000) (**Fig. 2—figure supplement 1A-I**) during the cuboidal-to-squamous transition of follicular epithelium. Therefore, these proteins likely redistribute or are substantially reduced from the cell junctions during the cuboidal-to-squamous transition. Since *α-tat* knockdown impaired cell-flattening (**Fig. 1C-D**), we next analyzed its effect on membrane protein trafficking during epithelial morphogenesis. We knocked down *α-tat* in somatic clones (*hs-flp, act>α-tat-IR*). Compared with controls (**Fig. 2A-D′, blue asterisk**), *α-tat* knockdown reduced cell size and increased cadherin localization at cell boundaries in both follicular (**Fig. 2B, B′** (red asterisk) and **Fig. 2—figure supplement 2A**) and MAG epithelia (**Fig. 2D, D′** (red asterisk) and **Fig. 2—figure supplement 2B**). Likewise, FasIII and Lgl accumulated at clone boundaries (*hs-flp, act>α-tat-IR*) (**Fig. 2—figure supplement 3A–H**).

**Figure 2.**
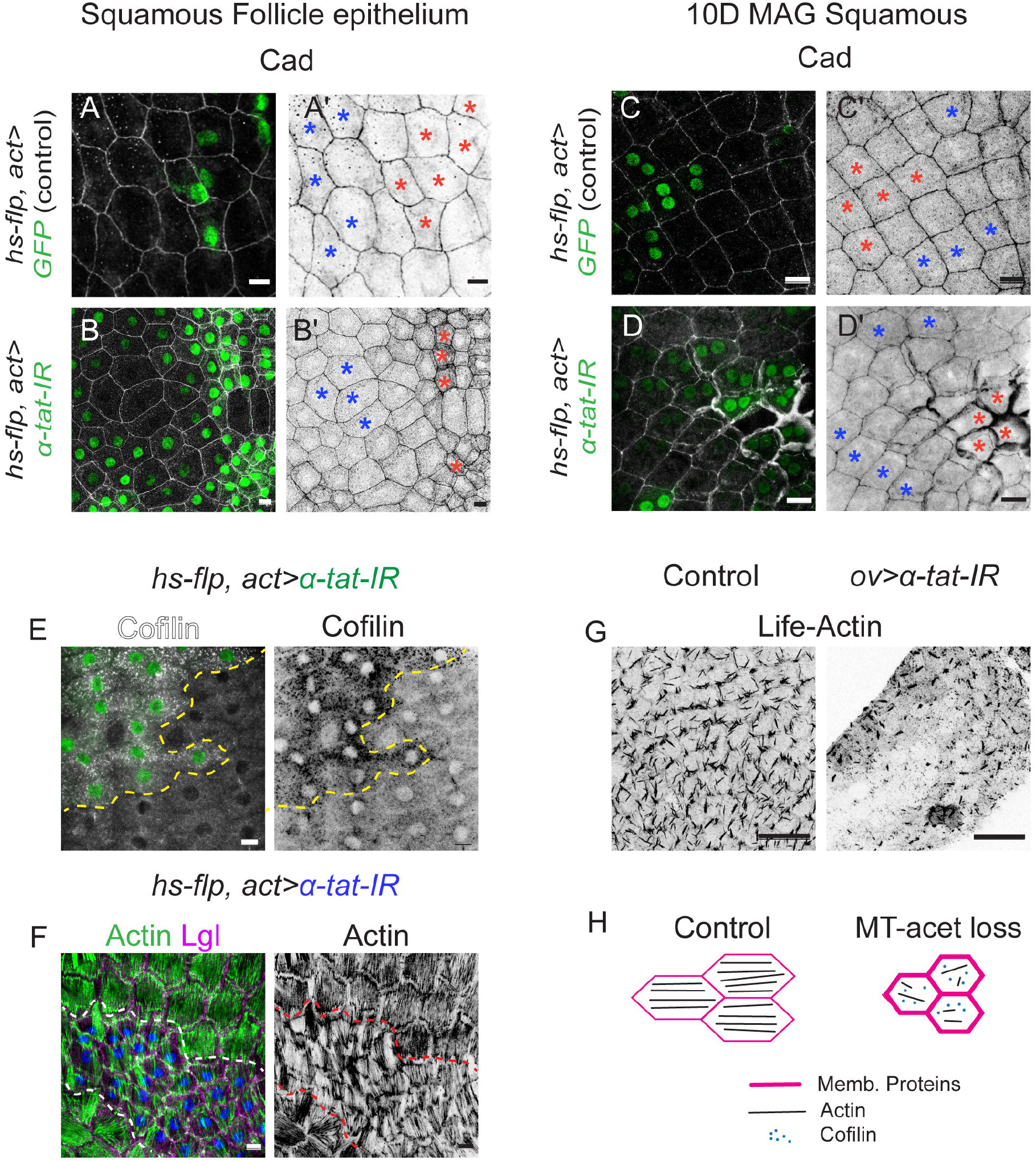
*α-tat* knockdown alters junctional assembly and actin organization in squamous epithelia. (**A, B**) Stage 14 ovarian follicular squamous epithelium displaying control (A, A’) and *α-tat* knockdown (B, B’) clones (marked by nuclear GFP). DE-Cad (*shg-mtomato* reporter) marks cell outlines. Compared to the neighboring wild type (blue star), DE-Cad levels in *α-tat-RNAi* cell boundaries are thicker (red stars). (**C, D**) Likewise, thickening of the Cadherin (DE-Cadherin antibody) at cell boundaries was also seen in *α-tat*-RNAi expressing somatic clones in MAG squamous epithelium from a 10-day-old adult. In both control (C) and test (D), somatic clones (nuclear GFP) are marked by red stars. (**E, F**) A comparison of cofilin (white/grey, E) and actin (phalloidin, green/grey, F) between control and somatic clones with *α-tat* knockdown (green nuclei, E and blue nuclei, F; also marked by broken lines) in the follicular epithelium of stage 14 oocytes. **(G)** Characteristics of Life-act GFP (grey) expression pattern in adult MAG epithelium and upon *α-tat* knockdown. **(H)** Schematic summarizing the cytoskeletal defects resulting from loss of microtubule acetylation. Scale bars: (A-F) 10 µm, (G) 50 µm

To further explore the consequences on vesicular trafficking, we analysed endosomal protein localization in *α-tat* knockdown somatic clones (*hs-flp, act>α-tat-IR*) in Stage 14 follicular epithelium. Sequestration of these membrane proteins was accompanied by peripheral accumulation of early endosomes (Rab5) containing Lgl (**Fig. 2—figure supplement 4A–C**), a phenotype resembling the defective Fasciclin 2 trafficking reported upon Tao kinase knockdown during the columnar-to-squamous transition (Gomez et al., 2012). Microtubule acetylation, therefore, appears critical for the endocytic trafficking of membrane proteins during epithelial cell flattening.

To further probe the role of microtubule acetylation on the actin cytoskeleton, we knocked down *α-tat* in MAG and ovarian follicular epithelium. In follicular epithelium, *α-tat* knockdown increased levels of the actin-severing protein cofilin (*hs-flp, act>α-tat-IR*, **Fig. 2E**), which likely explains their depolymerization of actin (phalloidin, **Fig. 2F**). Similarly, *α-tat* knockdown in the MAG epithelium led to sparse and depolymerized actin structures (*ov>UAS-Life-Actin RFP, α-tat-IR*, **Fig. 2G**). Thus, loss of microtubule acetylation impairs both membrane protein transport and actin polymerization, which are essential processes for cell flattening in squamous epithelia (**Fig. 2H**). In essence, microtubule acetylation is critical for cell flattening, likely by promoting endocytic trafficking and actin polymerization.

### Tissue-specific crosstalk between Yki signaling and microtubule acetylation

Microtubule hyperacetylation is a recognized prognostic marker in carcinomas, including SCC (Wattanathamsan and Pongrakhananon, 2023). We therefore hypothesized that Yki-mediated repression of TOR signaling restrains tumor-promoting microtubule acetylation. However, we noted that while in both follicular and MAG squamous epithelia, Yki exhibits nuclear localisation (Yki-GFP, **Fig. 3A–B**), the fallouts of its knockdowns were disparate. In the MAG, *yki* knockdown increased cell area (**Fig. 3—figure supplement 1A–C**, readout of enhanced TOR signaling (Bhattacharya et al., 2024)) with accompanying elevated microtubule acetylation (**Fig. 3C–E**), cytoarchitectural disruption (**Fig. 3F**), while *α-tat* knockdown reduced microtubule acetylation in *yki-*knockdown MAG (*ov>tat-IR, yki-IR;* **Fig. 3E**), and suppressed MAG-SCC (**Fig. 3F-G**) and extended host lifespan (**Fig. 3H**). By contrast, in the follicular epithelium, *yki* knockdown reduced cell size (**Fig. 3—figure supplement 1D–E)** and downregulated acetylated microtubules (**Fig. 3I–J**). As predicted by this model, gain of TOR signaling in the MAG (*ov>myr-Akt*, **Fig. 3K**) or follicular epithelium (*c204>myr-Akt*, **Fig. 3L**) increased microtubule acetylation. Conversely, TOR downregulation in the MAG (*ov>Akt-IR*, **Fig. 3M**) or follicle (*c204>Akt-IR*, **Fig. 3N**) decreased acetylation, confirming TOR as a direct regulator of microtubule acetylation *in vivo*.

**Figure 3.**
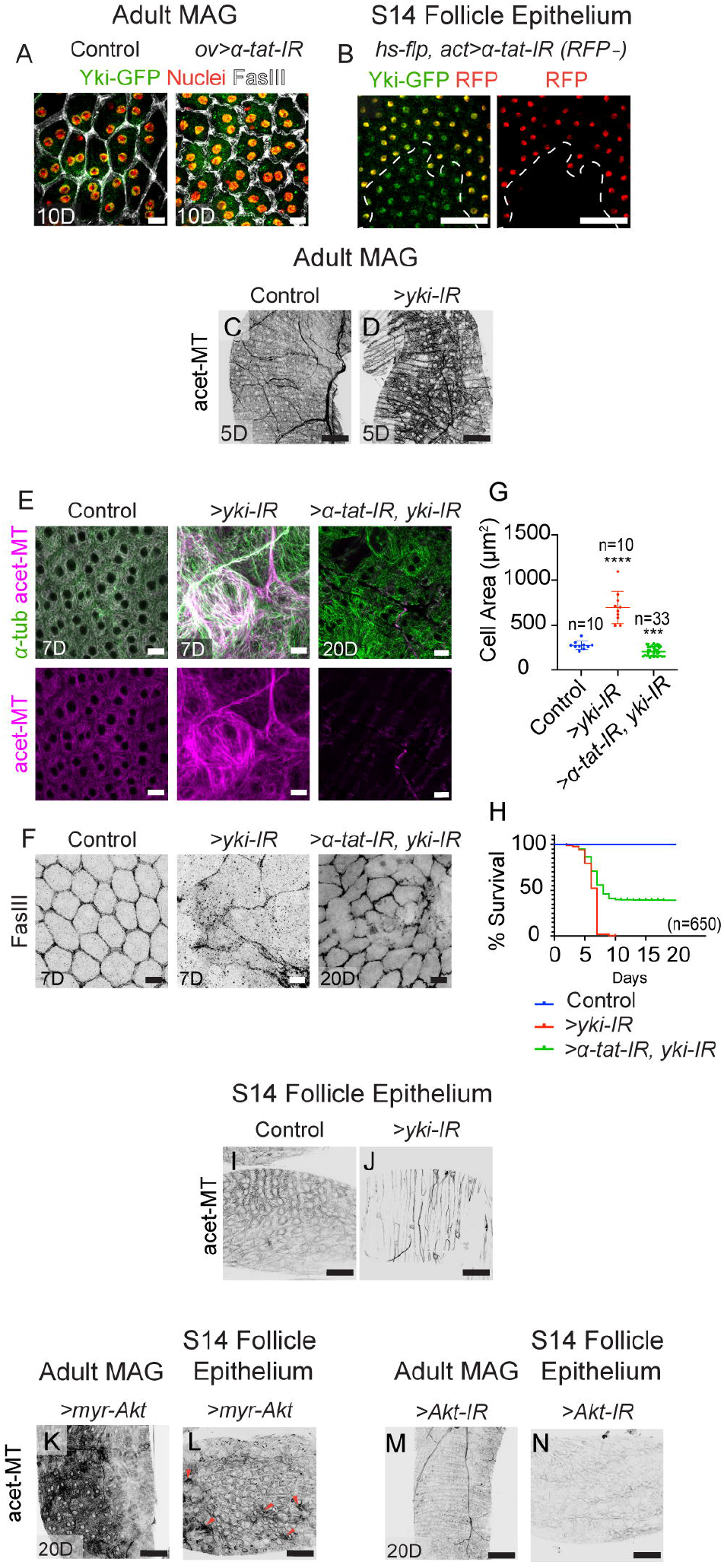
Yki linked TOR signaling regulate microtubule acetylation in squamous epithelia. (**A, B**) *α-tat* knockdown leads to persistent nuclear localization of Yki-GFP in both MAG (*ov>α-tat-IR*, A) and follicle (RFP negative cells, *hs-flp, act>a-tat-IR*, B) epithelia. (**C, D**) *yki* knockdown increases acetylated microtubule (acet-MT, grey) levels in the MAG epithelium (D) compared to control (C). (**E**) The elevation of acetylated microtubules caused by *yki* knockdown (*ov>yki-IR*, middle panel) is reversed by simultaneous *α-tat* knockdown *(ov>α-tat-IR; yki-IR*, right panel). (**F-H**) *yki* knockdown in the MAG epithelium increases cell area (F, G). Simultaneous knockdown of *yki* and *α-tat* restores MAG cell area (right panel, see quantification in G), besides adult survival (Kaplan-Meier plot, H). This phenotype and reduced adult survival (H) are rescued by co-knockdown of *α-tat*. (**I, J**) *yki* knockdown in the follicle epithelium (*GR1>yki-IR*) (J) reduces acetylated microtubule levels compared to control (I). (**K-N**) Gain of TOR signaling (via *myr-Akt*) elevates acetylated microtubules (acet-MT) in both MAG (*ov>myr-Akt*, K) and follicle (*c204>myr-Akt*, L) epithelia, while loss of TOR signaling (via *Akt-IR*) reduces it (*ov>Akt-IR*, M; *c204>Akt-IR*, N). Data analyzed by unpaired *t*-test (\*\*\*\**p<0*.*0001*, \*\*\**p=0*.*0008*). *n* values indicate cells; N=3 Scale bars: 10 µm (A, E, F); 50 µm (B-D, I-N).

The mechanism of TOR-dependent acetylation is likely conserved. We noted that mammalian ATAT1, the homolog of *Drosophila* α-tat, contains conserved AKT3- and pS6K-phosphorylation sites (**Fig. 3—figure supplement 2A**). Furthermore, in mammalian BS-C-1 cells, treatment with the TOR inhibitor rapamycin reduced microtubule acetylation (**Fig. 3— figure supplement 2B-C**). These findings suggest a conserved pathway for TOR-mediated regulation of α-tubulin acetylation.

Together, a Yki-dependent regulation of TOR signaling (and consequently microtubule acetylation), as seen in the MAG, shows a causal underpinning of the cell type-specific tumor suppressor role of Yki.

### A developmental priming of the Yki-TOR-microtubule acetylation axis

The absence of TOR-mediated microtubule hyperacetylation upon *yki* knockdown in follicular epithelium suggests tissue-specific developmental priming. To test this possibility, we analyzed single-cell transcriptomic data from 44,544 integrated cells of both tissues (Fly Cell Atlas). Out of twelve annotated cell clusters (**Fig. 4A**, see *Supporting Information*), we focused on MAG main cells and stage 14 choriogenic main-body follicle cells (**Fig. 4B**). Our analysis revealed that, compared to MAG squamous epithelia, follicular epithelium showed higher baseline expression of TOR-pathway genes (S6k, PDK1, Pten, Mitf, PI3K21B) and Insulin-pathway genes (rl, InR, foxo, chico, Ilp8, ImpL2) (**Fig. 4C**), consistent with their roles in supporting intense biosynthetic demands of chorion formation and oocyte growth (Das and Arur, 2017). In MAG, a relatively lower TOR/Insulin signaling may reflect evolutionary prioritization of nutrient allocation toward oogenesis over sex-peptide production. Follicular cells also exhibited elevated expression of the TEAD transcription factor Sd and its downstream effector Diap1 (**Fig. 4C**), indicating a Yki-linked pro-survival bias. We propose that these transcriptional differences arise from the distinct mechanostresses of each tissue (Gullekson et al., 2017; Hannezo and Heisenberg, 2019).

**Figure 4:**
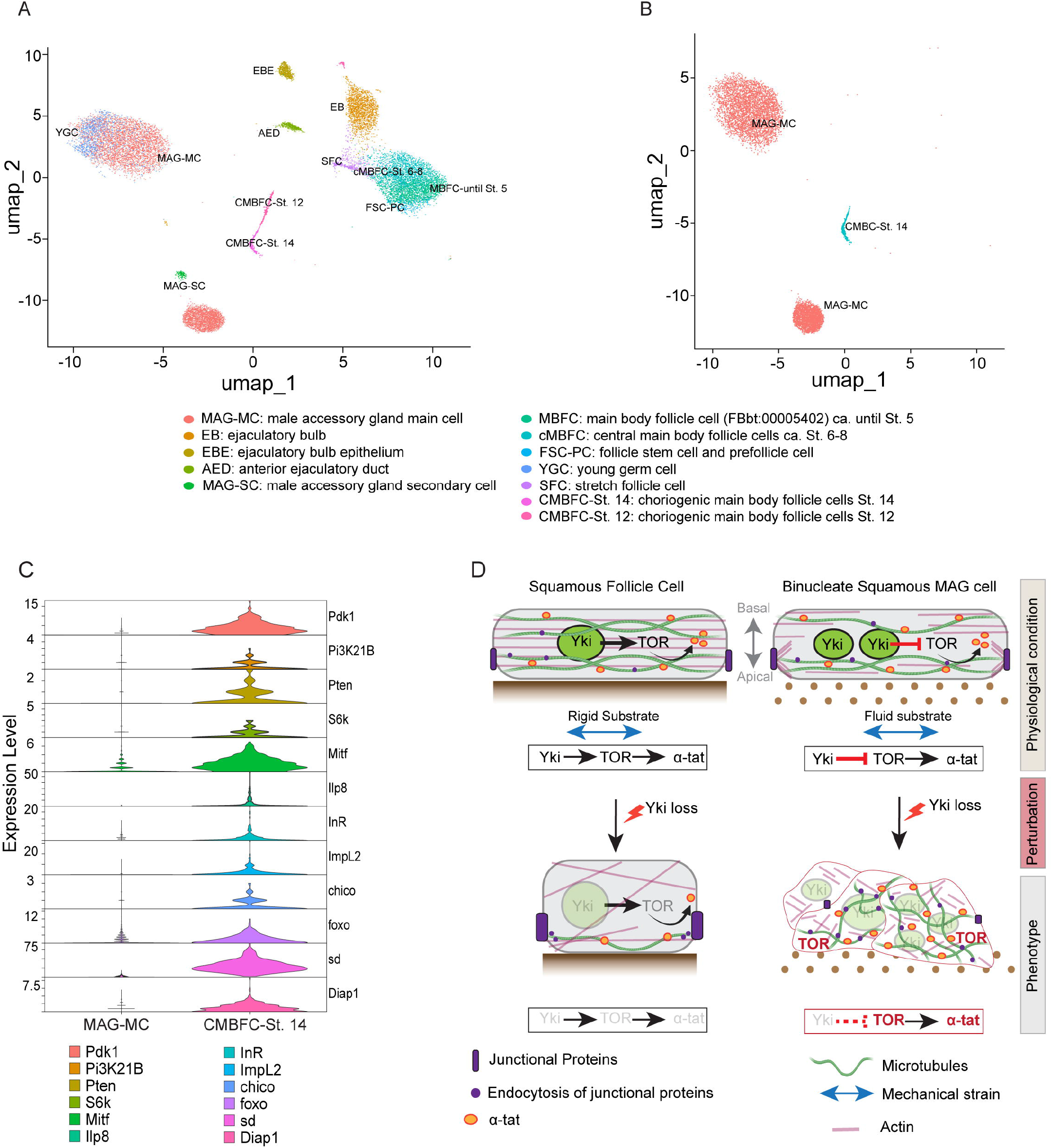
Distinct mechanical contexts drive differential Yki and TOR signaling in squamous epithelia of MAG and choriogenic follicular cells. **(A)** UMAP visualization of integrated single-cell transcriptomes from wild-type adult male accessory gland (MAG) and ovary. Cells are color-coded by their previously annotated cell types from the Fly Cell Atlas. **(B)** UMAP of a subset containing only the squamous epithelial cells: choriogenic main body follicle cells at Stage 14 (CMBFC-St.14) from the ovary and main cells from the MAG (MAG-MC). This subset was used for comparative analysis of squamous epithelia from male and female reproductive organs. **(C)** Violin plots showing the expression levels of key genes from the TOR, Insulin, and Hippo signaling pathways, identified through differential expression analysis between MAG-MC and CMBFC-St.14 cells. **(D)** Model of mechanical context-dependent Yki signaling outputs. The ovarian follicular squamous epithelium (left) interfaces with a rigid apical chorion, while the MAG squamous epithelium (right) faces a fluid-filled lumen. Knockdowns of the core mechanosensing component, nuclear Yki signaling, lead to distinct phenotypes in these two mechanical contexts: a squamous-to-cuboidal transition in the follicle epithelium, but squamous cell carcinoma (SCC) in the MAG.

## Discussion

Protooncogenes like YAP/TAZ/Yki can function as context-dependent tumor suppressors (for review, see (Baroja et al., 2024; Franklin et al., 2023; Luo et al., 2023)), or “double agents” (Shen et al., 2018) but the mechanisms behind this duality have remained elusive. Here, we identify the suppression of TOR-driven α-tubulin acetylation as the mechanistic basis for Yorkie’s (Yki) tumor suppressor role in the *Drosophila* male accessory gland (MAG) squamous epithelium.

Variations in substrate stiffness affect mechanochemical sensing (Hannezo and Heisenberg, 2019). Tissue-specific outcomes of Yki loss seen here likely mirror this mechanochemical distinctiveness of MAG and follicular squamous epithelium. Previously, it was also shown that higher substrate stiffness enhances α-tubulin acetylation, actin bundling, and YAP/TAZ nuclear entry (Seetharaman et al., 2022). While both MAG and follicular epithelia exhibit constitutive nuclear Yki, only the MAG develops carcinoma upon Yki loss, suggesting that their disparate substrate stiffness may underlie this differential tumor susceptibility. This interpretation is corroborated by the distinctiveness between the two tissues. First, the MAG epithelium faces a fluid-filled lumen, whereas the follicular epithelium interfaces with a rigid chorion enveloping the oocyte. Second, our single-cell transcriptomics revealed that follicular cells are developmentally primed with high baseline TOR/Insulin signaling and Yki-target gene expression, likely fulfilling the biosynthetic demands of oogenesis. In contrast, MAG epithelium relies on Yki to actively restrain the TOR-α-tubulin acetylation axis, making it uniquely susceptible to Yki-loss-induced SCC (**Fig. 4D**).

Our findings integrate several isolated pillars of human SCC pathogenesis. For instance, in alveolar type 1 (AT1) lung cells, nuclear YAP/TAZ activity restrains TOR signaling and maintains cell homeostasis (Ohnishi et al., 2024), oncogenic PI3K-Akt-TOR signaling is a known driver of SCCs (Zheng et al., 2024), and α-tubulin acetylation promotes SCC aggressiveness and chemoresistance (Wattanathamsan and Pongrakhananon, 2023). Our findings provide a unified, causal pathway linking these elements: the loss of a Yki signaling “double agent” triggers TOR signaling, which in turn drives microtubule hyperacetylation to cause carcinoma. By delineating the Yki-TOR-α-tubulin acetylation axis, we provide a mechanistic model for how “double-agent” protooncogenes can suppress cancer in specific tissues, revealing new potential avenues for therapeutic intervention in squamous cell carcinomas.

## MATERIALS AND METHODS

All the genetic stocks and reagents used in this study have been procured from public repositories or as gifts from other researchers. Details of the experiments, for instance, *Drosophila* maintenance and induction of transgenes, immunostaining protocol, mammalian cell culture-based protocol, microscopy, quantification, statistical analysis, bioinformatics, and scRNA sequence analysis, are provided in the *Supporting Information Appendix*.

## DATA AVAILABILITY

All data generated during this study are included in this article. Additionally, publicly available data from the Fly Cell Atlas were analyzed in this research.

## ACKNOWLEDGEMENTS

We thank Barry Thompson, Kristipati Ravi Ram, and Mohit Prasad for fly stocks, and Jonaki Sen for the cofilin antibody. We also thank Debdeep Dutta, Thamarailingam Athilingam, and Saurabh Singh Parihar for their critical manuscript reading.

