## Supporting Information for "Paradoxical Tumor Suppressor Role of Yorkie Through a TOR‑Dependent α‑Tubulin Acetylation in Select Squamous Epithelium"

*Title*

*Running title*

**Lineage-specific tumor suppressor role of Yki**

*Authors*

Rachita Bhattacharya[1, 4], Jeganath Ammavasai[1], Shruti Agarwal[1, 2], Hamim Zafar[1, 2, 3], Pradip Sinha[1, *], and Nitin Mohan[1, *]

[1]Biological Sciences and Bioengineering, [2]Mehta Family Center for Engineering in Medicine, [3]Computer Science and Engineering, Indian Institute of Technology Kanpur, India
[4]Current address: Genetics and Developmental Biology Unit, Institut Curie, CNRS UMR 3215, INSERM U934, Paris, France

*Corresponding authors: Pradip Sinha and Nitin Mohan

**Author Contributions:** Conceptualization: RB, NM, PS. Data acquisition: RB, JA. Transcriptomic analysis: RB, SA, HZ. Manuscript draft, edits, and revision: RB, NM, PS.
**Competing Interest Statement:** The authors declare no conflict of interest.
**Classification:** Biological Sciences, Developmental Biology
**Keywords:** Yorkie, TOR, microtubule acetylation, squamous cell carcinoma.

**This file includes:**

1. Experimental model and method details
2. References
3. Supplementary figure legends

### EXPERIMENTAL MODEL AND METHOD DETAILS

#### *Drosophila* maintenance and targeted gene expression

*Drosophila* cultures were maintained at 25°C on standard yeast cornmeal agar food. The genetic knockdown and overexpression experiments in MAG and follicles were done at 29°C to increase the efficiency of Gal4 activity (1). The *ov-Gal4* (2) driven expressions of desired transgenes on the MAG were performed as described previously(3). For studies using the *GR1-Gal4* (4) and *c204-Gal4* drivers (5), freshly eclosed female flies were shifted to 29°C and raised on yeast-paste supplement food for 4-5 days until dissection.

We examined squamous epithelial morphology in MAG post-epithelial flattening in four-day-old males (3) and stage 14 of the egg chamber of ovarian follicles (6). For the induction of a desired gene in somatic clones, we used the flp/FRT-flip-out technique (7). For crosses involving *hs-flp, act>CD2>Gal4, UAS-GFP*, a heat shock at 37°C was applied to 5-day-old adults—5 minutes for males and 10–20 minutes for females. For crosses with *hs-flp, act>CD2>Gal4 UAS-stinger*, a 30-minute heat shock was given to adult females 3 times a day. Following heat shock, females were shifted to 29°C and fattened on yeast-paste for 5 days until dissection (8), while males were dissected 7–10 days after heat shock.

The details of the fly stocks, reagents, and resources are as follows:

*Fly stocks:*

*UAS-GFP.nls* (BDSC#4776), *UAS-yki-RNAi* (BDSC#34067), *UAS-myr-Akt* (BDSC#80935), *UAS-Akt-RNAi* (BDSC#82957), *FasIII-GFP* (BDSC#59809), *shg-mtomato* (BDSC#58789), *UAS-α-tat-RNAi* (9) (Chr. II, BDSC#62331), *UAS-α-tat-RNAi* (Chr. III, BDSC#28777), *UAS-Lifeact-RFP* (BDSC#58362), *GR1-Gal4* (BDSC#36287), *c204-Gal4* (BDSC#3751), *hs-flp, act>CD2>Gal4, UAS GFP (nls)* (Lab generated), *hs-flp, act>CD2>Gal4, UAS stinger (nls)* (Gift from Mohit Prasad), *Yki-GFP* (Gift from Barry Thompson), *ovulin-Gal4* (Gift from Kristipati Ravi Ram).

*Reagents:*

Mouse anti-FasIII (DSHB, 7G10), Mouse anti-α-tubulin (DSHB, 12G10), Mouse anti-Cofilin (Santa Cruz, Gift from Jonaki Sen), Rabbit anti-α-tubulin (Abcam, ab18251), Mouse anti-acetylated tubulin (Sigma-Aldrich, T6793), Alexa Fluor 488 Goat anti-mouse IgG (Invitrogen, A11007), Alexa Fluor 555 Goat anti-mouse IgG (Invitrogen, A32727), Alexa

Fluor 633 Goat anti-mouse IgG (Invitrogen, A21050), Alexa Fluor 488 Goat anti-rabbit IgG (Invitrogen, A21206), Alexa Fluor 555 Goat anti-rabbit IgG (Invitrogen, A21428), Alexa Fluor 633 Goat anti-rabbit IgG (Invitrogen, A21070), TO-PRO-3 (Invitrogen, S33025), Phalloidin-555 (Invitrogen, A34055), Vectashield (Vector Laboratories, #H-1000).

*Software and Algorithms:*

Leica LAS AF software (Leica microsystem), Prism (Graphpad), Illustrator (Adobe CC), FIJI (ImageJ), RStudio (Seurat version 4.0 (10))

**Immunostaining and microscopy**

MAG and ovarian follicles were dissected in cold PBS and fixed in 4% paraformaldehyde. Tissues were permeabilised in PBST (1XPBS containing 0.02% Triton-X-100). After blocking in 5% BSA for 2 h, they were incubated with primary antibody overnight at 4°C, washed three times for 10 min each in PBST, and then incubated with appropriate fluorescent-tagged secondary antibodies (1:250, Invitrogen) for two hours at room temperature. The samples were washed three times for 10 min each in PBST, followed by two washes for 10 min each in PBS, and counterstained with TO-PRO-3 and/or Alexa Fluor™ 555 phalloidin (A34055, Invitrogen), wherever appropriate. Finally, the tissues were washed thrice for 10 min each in PBS, mounted in Vectashield®, and sealed with nail paint. The samples were imaged with a Leica SP5 or Stellaris laser scanning confocal microscope. All images were assembled using Adobe Illustrator.

**Quantification and statistical analysis**

Cell area measurements were used as a proxy for columnar-to-squamous transition, as done in previous studies (11, 12). Larger cell areas indicated squamous or flattened cells, while smaller areas corresponded to columnar cell types (11, 12). All cell areas and fluorescence intensity were calculated using FIJI. A freehand drawing tool was used to trace the cell or organ boundaries and calculate the surface area and integrated density. All image acquisitions, particularly those involving fluorescence intensity measurements, were performed using the same parameters for control and test samples. To quantify the fluorescence intensity of acetylated tubulin in the columnar and squamous follicle areas, the integrated density in each case was normalised to its respective average cell area. The quantification of percentage acetylated microtubules was done by calculating the acetylated tubulin fluorescence intensity levels relative to total α-tubulin (13) for columnar and squamous follicle epithelia at Stage 10 and Stage 14, respectively. The membrane protein intensity during follicle cell stretching was quantified by normalizing the integrated density in columnar and squamous epithelia to the maximum intensity across both cell types. To compare the fluorescence levels of Cad, FasIII, and Lgl in somatic clones of *α-tat-RNAi* with the control, the ratio of the integrated density of membrane proteins in clone vs. non-clone cells was calculated for both the control and *α-tat* knockdown conditions and graphically plotted. The colocalisation of Rab5 vesicles with Lgl was calculated using the plugin Coloc 2 in FIJI (https://imagej.net/Coloc_2).

All statistical analyses were performed using GraphPad 9.0. An unpaired *t*-test was performed to quantify cell, organ, and nuclear sizes. A cutoff of $p$>0.05 was used to define statistical significance in all graphical plots. The values of n (number of cells) and N (number of samples) for each experiment are provided in the figures or figure legends, with phenotype frequencies indicated in the relevant images. Illustrations were made using Adobe Illustrator.

**Single Cell RNA Sequencing Analysis**

Raw data for *Drosophila* males and females were obtained from Fly Cell Atlas (14) in h5ad format, which was already annotated. The counts from both the male dataset (13143 cells) and the female dataset (31401 cells) were extracted and concatenated into a single Anndata object using Scanpy (15). Batch integration of the concatenated data was performed using scDREAMER (16), which infers cellular latent embeddings. Subsequently, the Anndata object was converted into a Seurat (17) object using the 'schard' package and 'h5ad2seurat' function in R. We incorporated scDREAMER embeddings as reductions of the Seurat object using 'CreateDimReducObject'. For visualizing the cells, the scDREAMER (16) embeddings of the cells were projected in two dimensions using the Uniform Manifold Approximation and Projection (UMAP) algorithm (18). The 'FindNeighbors' and 'FindClusters' functions in Seurat were used for clustering the cells based on scDREAMER embeddings. The resulting cell clusters were visualized in UMAP space using the 'DimPlot' function in Seurat. The expression of marker genes by cells in the clusters was visualized using the 'FeaturePlot' and 'VlnPlot' functions. The differentially expressed genes in each cluster were inferred using the 'FindAllMarkers' function in Seurat.

For further analysis, the 'Subset' function of Seurat was used to subset male accessory gland main cells and choriogenic main body follicle cells at stage 14 to create a Seurat object containing 6919 cells. Differential gene expression (DGE) analysis was performed between the two squamous epithelia using the 'FindMarkers' function of Seurat, and significant DEGs were used to prepare violin plots.

**Cell culture, transfection, and drug treatment.**

African green monkey (*Chlorocebus pygerythrus*) kidney cells (BS-C-1; ATCC) were maintained in complete growth medium (minimum essential medium supplemented with non-essential amino acids, 10% FBS, 2 mM L-glutamine, 1 mM sodium pyruvate, and penicillin-streptomycin) at 37°C with 5% $CO_2$.

Plasmids encoding Raptor-GFP, myc-mTOR, and mlst8-myc (Addgene) were propagated in bacterial cultures and purified using Qiagen or Favorgen kits. Plasmid identity was confirmed by restriction digestion. BS-C-1 cells were seeded in 8-well Lab-Tek chambered coverglass or on confocal imaging discs 12–24 hours prior to transfection and maintained in complete growth medium. Triple transfection was performed using the Effectene Transfection Kit (Roche) according to the manufacturer's instructions, with plasmids mixed in a 1:2:2 mass ratio (Raptor-GFP:myc-mTOR:mlst8-myc). Cells were incubated for 24 hours post-transfection at 37°C in 5% $CO_2$ before imaging or fixation.

For Rapamycin treatment, cells were incubated with 0.2 μM Rapamycin (Merck, R0395) for 1 hour (19).

**Immunofluorescence.** Cells were seeded on 8-well Lab-Tek chambered coverslips and fixed with 3% PFA and 1% GA for 10 min at room temperature. After blocking with 3% BSA and 0.2% TritonX-100, cells were incubated with primary antibodies for 1 hour at room temperature or overnight at 4°C. Following three washes with 0.2% BSA and 0.05% TritonX-100, cells were incubated with secondary antibodies for 40–60 min and washed thrice with PBS. For double staining, primary or secondary antibodies were incubated simultaneously. Imaging was performed on a Nikon Ti2E microscope with TIRF (100X oil immersion objective) using 488, 560, and 633 nm lasers.

**Super-resolution Structured Illumination Microscopy (SIM).** SIM was performed using a ZEISS Elyra 7 system (Lattice SIM) with Plan-Apochromat 63x/1.4 oil immersion objective and appropriate filter cubes. Excitation wavelengths were 488, 561, and 642 nm. Optical sectioning (Z-stacks) was collected in 3D. Image processing used the manufacturer's software in 3D standard mode.

**SIM image processing and MT acetylation quantification.** SIM image channels (α-tubulin and acetylated α-tubulin) were maximum or sum intensity projected along the z-axis, converted to 16-bit images, and combined to capture total microtubule (MT) populations. Images were auto-thresholded using the Isodata algorithm in FIJI. The Mander's coefficient (overlap fraction) of acetylated MT over total MT was computed using the JACoP plugin and multiplied by 100 for percentage acetylation.

**Phosphorylation site analysis.** To explore potential regulation of microtubule acetylation by mTORC1 signaling kinases (AKT, mTOR, S6K1), phosphorylation sites on α-tubulin acetyltransferase were identified using the PhosphoSite Plus tool (20).

**Supplementary Figure legends:**

**Figure 1—figure supplement 1. Cell flattening and microtubule acetylation are correlated and *α-tat* dependent**.

(**A, B**) Quantification of cell area in follicular (A) and MAG (B) epithelia reveals the transition to squamous morphology.

(**C, D**) Acetylated α-tubulin levels are significantly higher in squamous epithelia of both the follicle (C) and the MAG (D).

(**E, F**) Knockdown of *α-tat* significantly reduces cell area in follicle (E) and MAG (F) squamous epithelia.

Data are mean ± SD; analyzed by unpaired two-sided t-test (****p<0.0001, **p=0.0005). n indicates number of cells.

Scale bars: 10 μm.

**Figure 1—figure supplement 2. Microtubule acetylation increases with total microtubule levels during cell flattening.**

(**A**, **B**) Representative images of total α-tubulin and acetylated α-tubulin in Stage 10 (columnar) and Stage 14 (squamous) follicle epithelia.

(**C, D**) Acetylated α-tubulin levels correlate with total α-tubulin in both columnar (C, $p$=0.0002) and squamous (D, $p$=0.02) stages.

(**E**) Quantifying the percentage of acetylated α-tubulin (acet-MT/total α-tub) in Stage 10 vs Stage 14 epithelia; ns=not significant

Data are mean ± SD; analyzed by unpaired *t*-test. N indicates sample number.

Scale bars: 10 μm.

**Figure 1—figure supplement 3. *α-tat* depletion reduces cell size without altering microtubule organization.**

(**A**, **B**) *α-tat* knockdown in somatic clones (*hs-flp, act*>*α-tat-IR*, blue nuclei, dashed outlines) in follicle (A) and MAG (B) epithelia show loss of acetylated α-tubulin but normal microtubule organization.

(**C, D**) Compared with the control stage 14 oocyte (C), *α-tat* knockdown in developing follicle epithelium (*GR1-Gal4*>*α-tat-IR*) reduces cell and oocyte sizes.

Scale bars: (A, B) 10 μm; (C, D) 100 μm.

**Figure 2—figure supplement 1**. **Junctional protein levels decrease during the transition to squamous morphology**.

(**A-I**) Immunostaining and quantification to reveal dilution of FasIII (A-C), DE-Cadherin (D-F), and Lgl (G-I) during the flattening of follicle epithelium from Stage 10 (columnar epithelium, CE) to Stage 12 (squamous epithelium, SE). Boxes (broken line) are shown at a higher magnification in respective insets.

Data are mean ± SD; analyzed by unpaired *t*-test (**p<0.0001*). *n*=number of cells; N=2.

Scale bars: main panels, 50 µm; insets, 10 µm

**Figure 2—figure supplement 2. *α-tat* knockdown increases junctional Cadherin levels**.

(**A**, **B**) Quantification confirms a significant increase in normalized Cadherin intensity at the membrane (*shg-mtomato*, A; DE-cadherin antibody, B) upon *α-tat* knockdown in both follicle (A) and MAG (B) squamous epithelia.

Data are mean ± SD; analyzed by unpaired two-sided *t*-test *(****p<0.0001, ***p=0.0003*). n values indicate number of cells; N=3.

**Figure 2—figure supplement 3. *α-tat* knockdown impairs the dilution of FasIII and Lgl**.

(**A-B**) *α-tat* knockdown clones (red stars) retain higher levels of the membrane proteins FasIII in both follicle (A, B) and MAG squamous epithelia (C, D). Likewise, these *α-tat* knockdown clones (red stars) display accumulation of Lgl at cell boundaries of both follicle and MAG (red star, E, F), compared to wild-type neighbors (marked by blue stars). Quantification of normalized fluorescence intensity for FasIII (B, D) and Lgl (F, H).

Data are mean ± SD; analyzed by unpaired t-test (*****p<0.0001; ***p=0.0003*). *n* values indicate cells; N=3.

Scale bar: 10 µm.

**Figure 2—figure supplement 4. Loss of microtubule acetylation disrupts early endosomal trafficking is squamous follicular epithelium**.

(**A**, **B**) *α-tat* knockdown clones (blue nuclei) show Rab5-positive early endosomes (green) accumulating at the cell periphery, colocalized with Lgl (magenta, yellow arrowheads). Quantification confirms increased Lgl-Rab5 colocalization (B).

(**C**) Schematic model: Loss of acetylated microtubules sequesters membrane proteins like Lgl with early endocytic vesicles.

Data are mean ± SD; analyzed by unpaired t-test (p=0.0002). *n* values indicate cells; N=3.

Scale bars: (A) 50 µm; (A' insets) 10 µm

**Figure 3—figure supplement 1. *yki* knockdown has opposing effects on cell area in MAG and follicle epithelia**.

(**A-C**) *yki* knockdown in the MAG epithelium (B) increases cell area (a hallmark of TOR activation) compared to control (A), as quantified in (C).

(**D**, **E**) In contrast, *yki* knockdown in the Stage 14 follicle epithelium (D, E) decreases cell area compared to control.

Data are mean ± SD; analyzed by unpaired *t*-test (*$p<0.0001$). *n* values indicate cells; N=3.

Scale bar: 10 µm

**Figure 3—figure supplement 2**. **mTOR signaling regulates microtubule acetylation in mammalian cells**.

(**A**) Phosphorylation site prediction for the ATAT1 enzyme (α-tubulin acetyltransferase) from the Phosphosite Plus database.

(**B**) BS-C-1 WT cells are untreated or transiently transfected with hyperactive mutant mTORC1 plasmids or treated with rapamycin (0.2 µM, 1hr) and stained against alpha (green) and acetylated microtubule (magenta), the inset image represents hyperactive mutant mTORC1 expression.

(**C**) Bar plot shows the percentage of acetylated microtubules under the mentioned conditions.

Data analyzed by one-way ANOVA followed by Tukey's test and presented as mean ±SD: ****$p<0.0001$.

Scale bar: 10 µm.
