## Supplementary figures and images for "Paradoxical Tumor Suppressor Role of Yorkie Through a TOR‑Dependent α‑Tubulin Acetylation in Select Squamous Epithelium"

### Figure 1-figure supplement 1

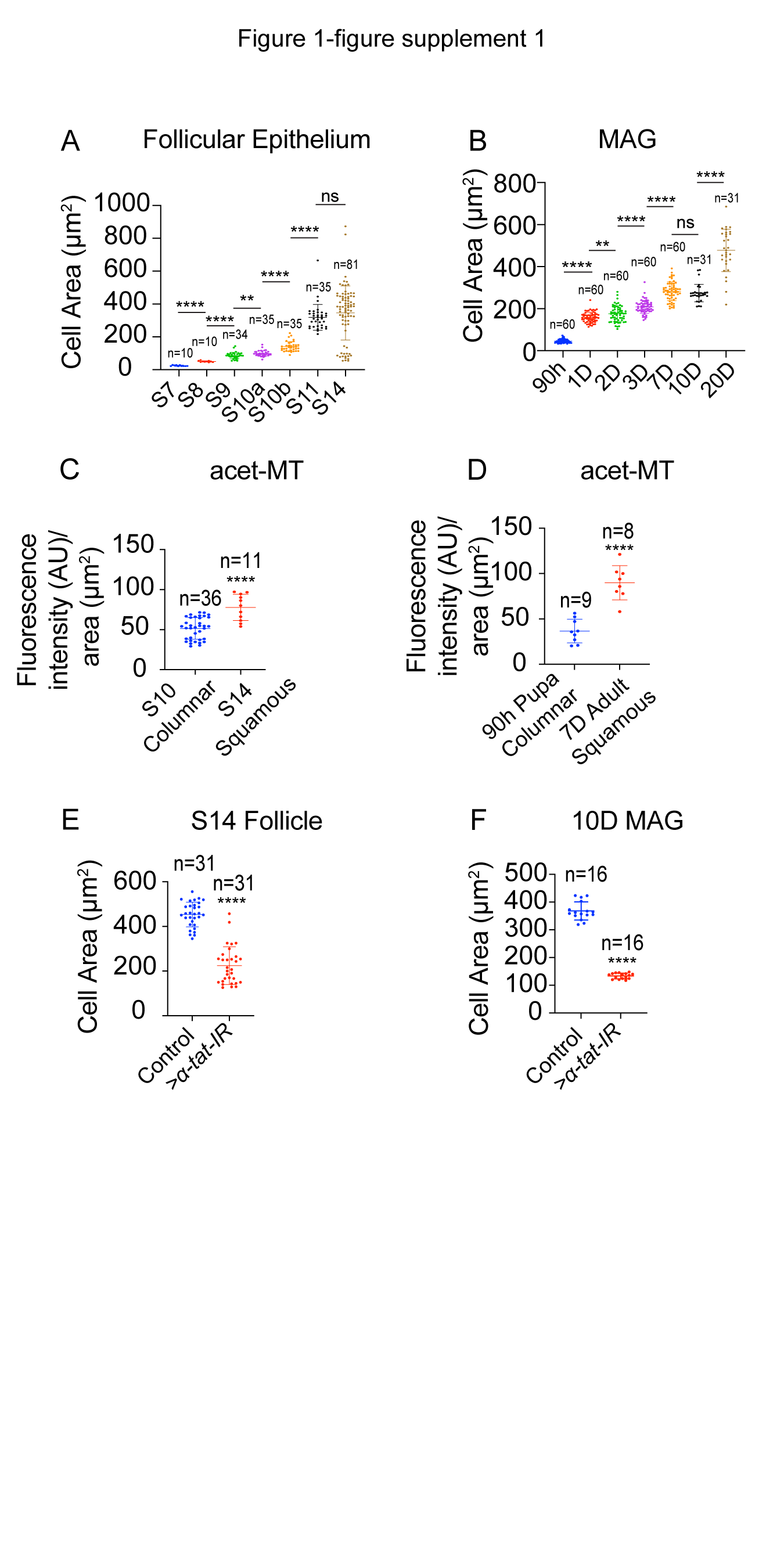

### Figure 1-figure supplement 2

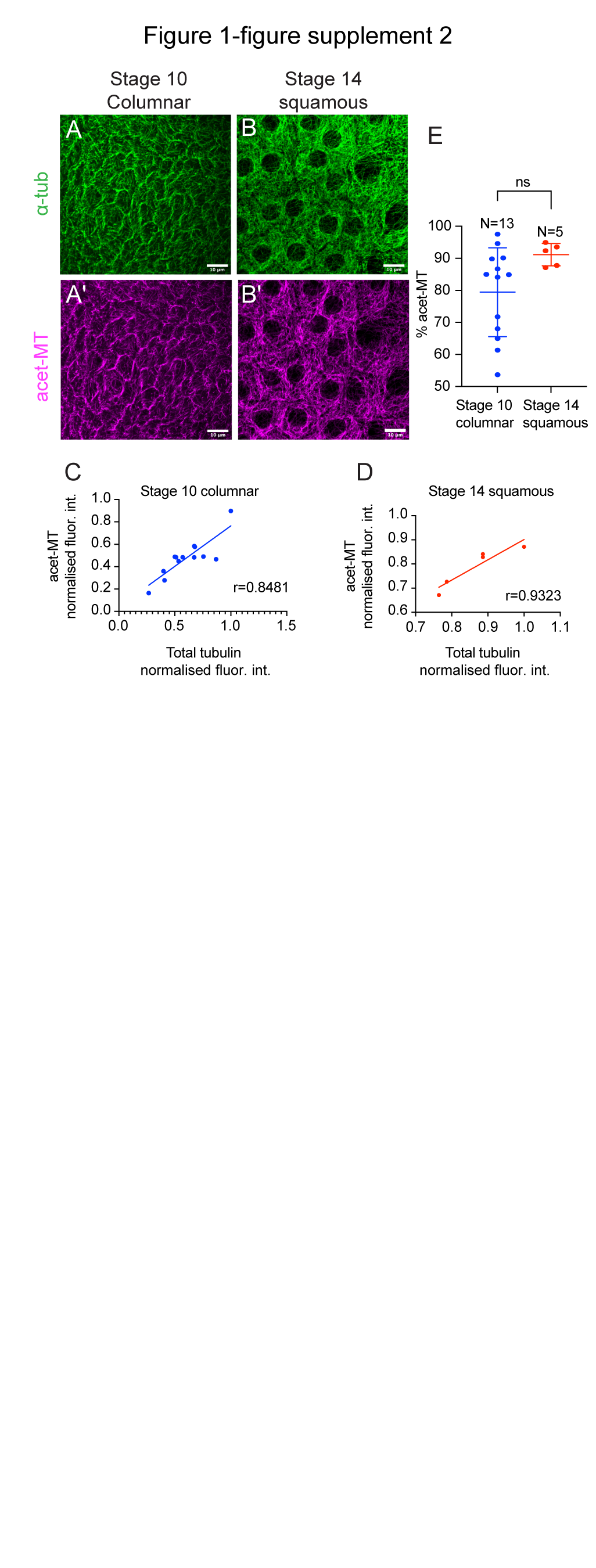

### Figure 1-figure supplement 3

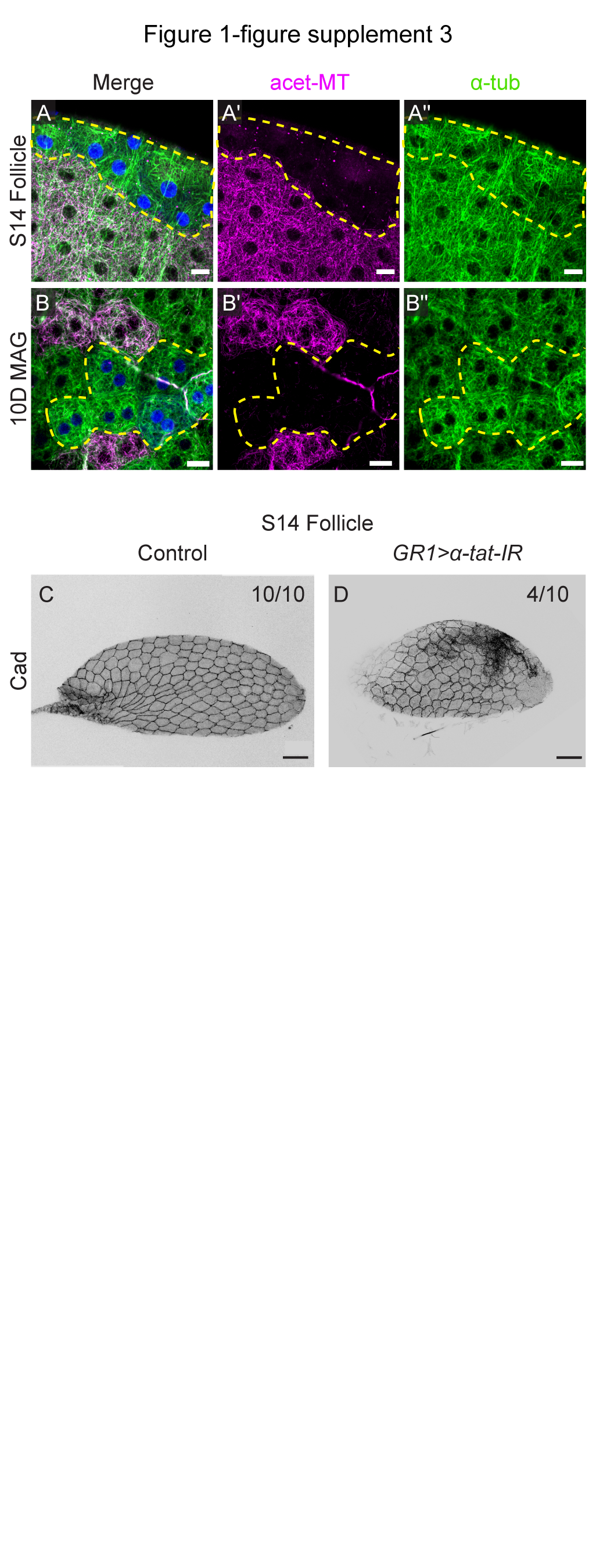

### Figure 2-figure supplement 1

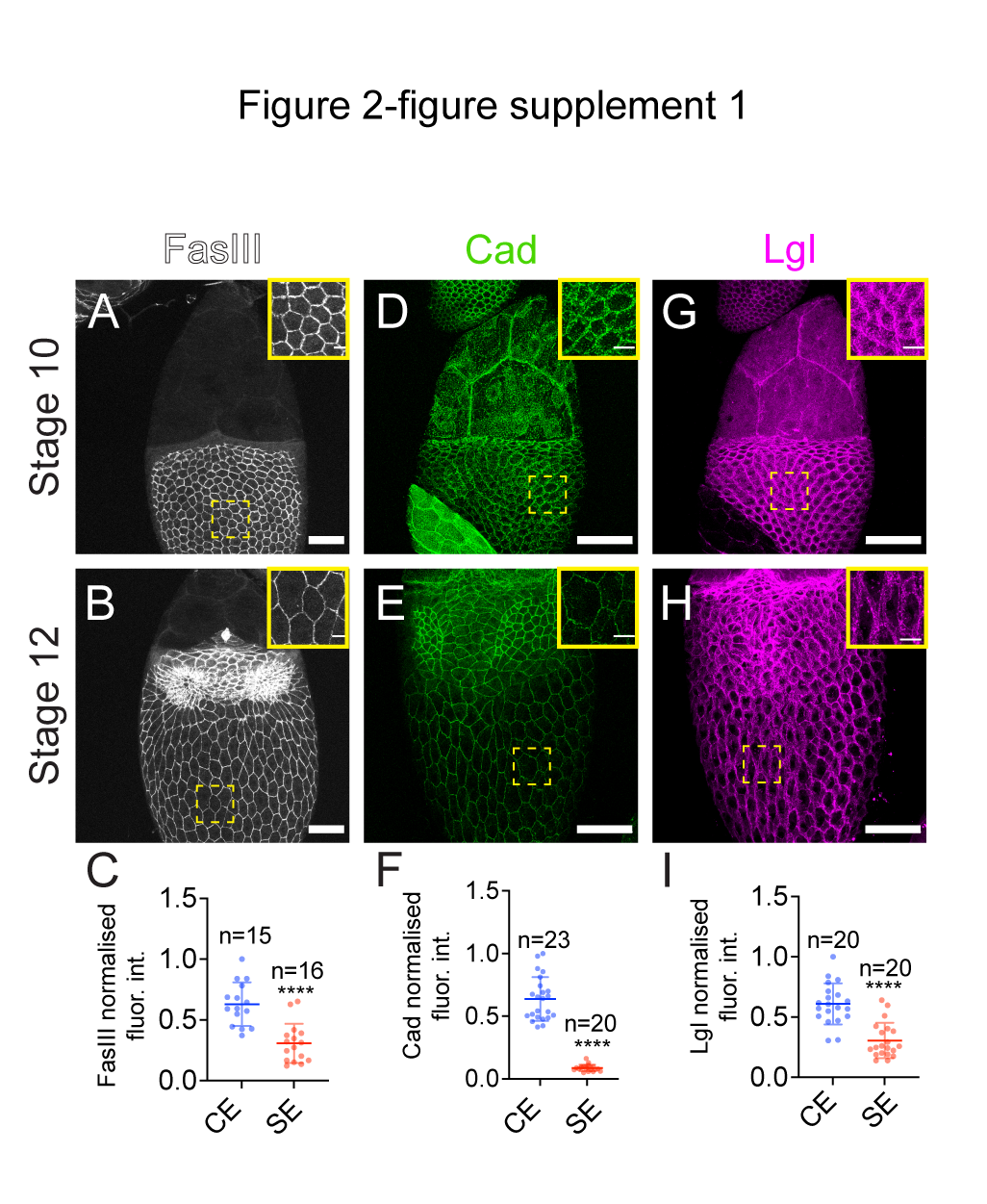

### Figure 2-figure supplement 2

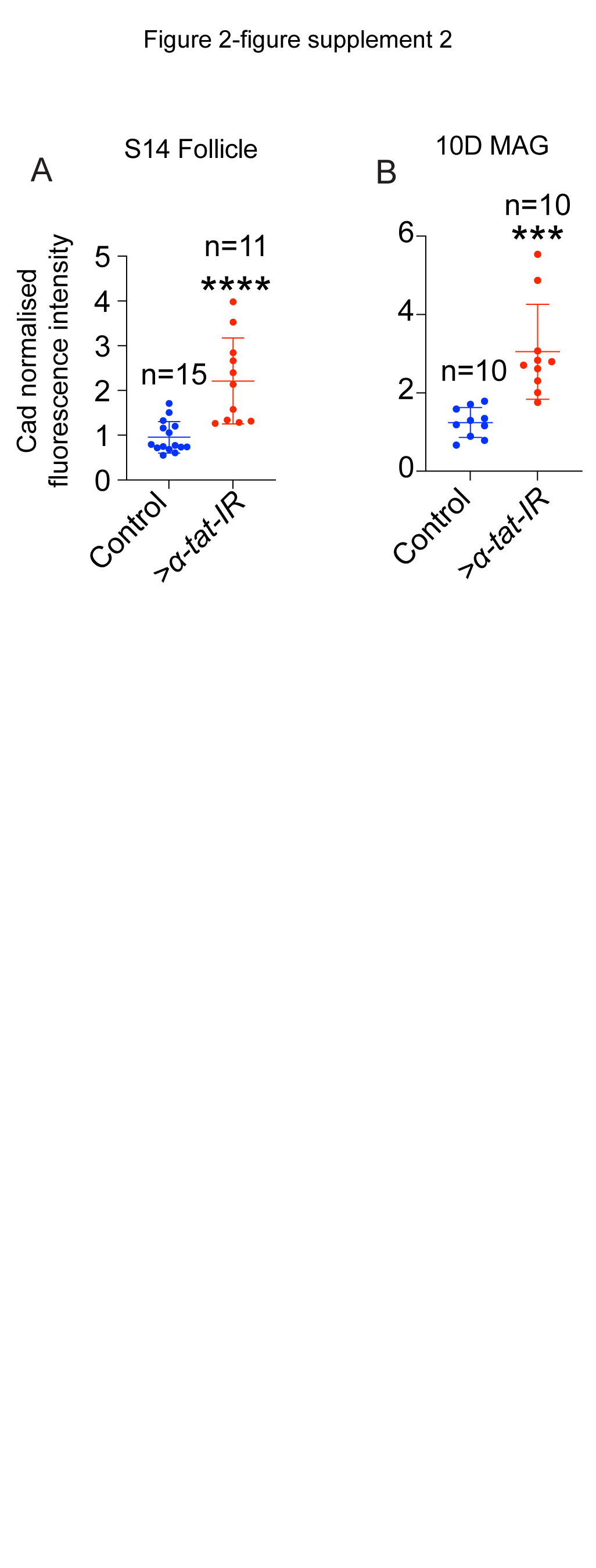

### Figure 2-figure supplement 3

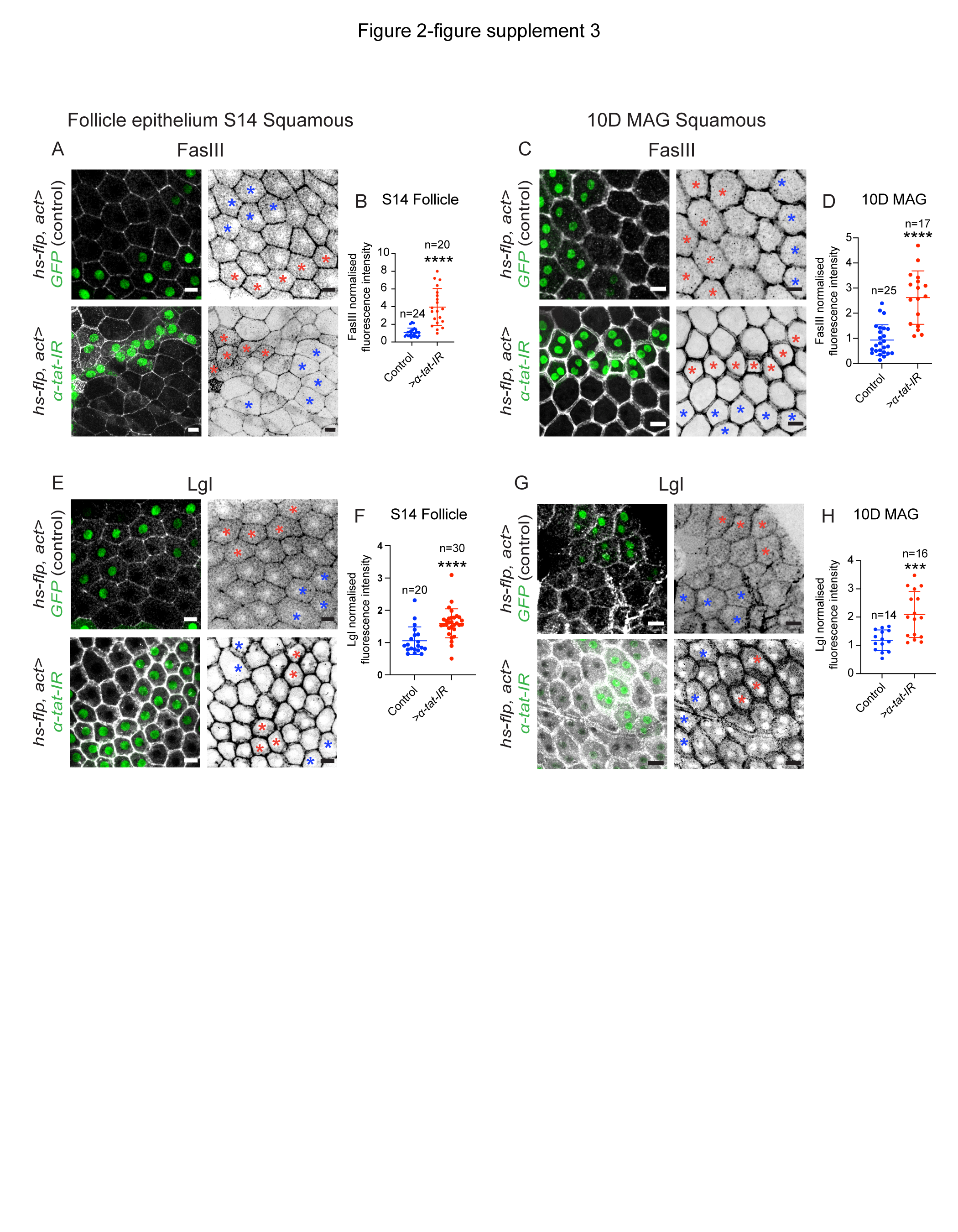

### Figure 2-figure supplement 4

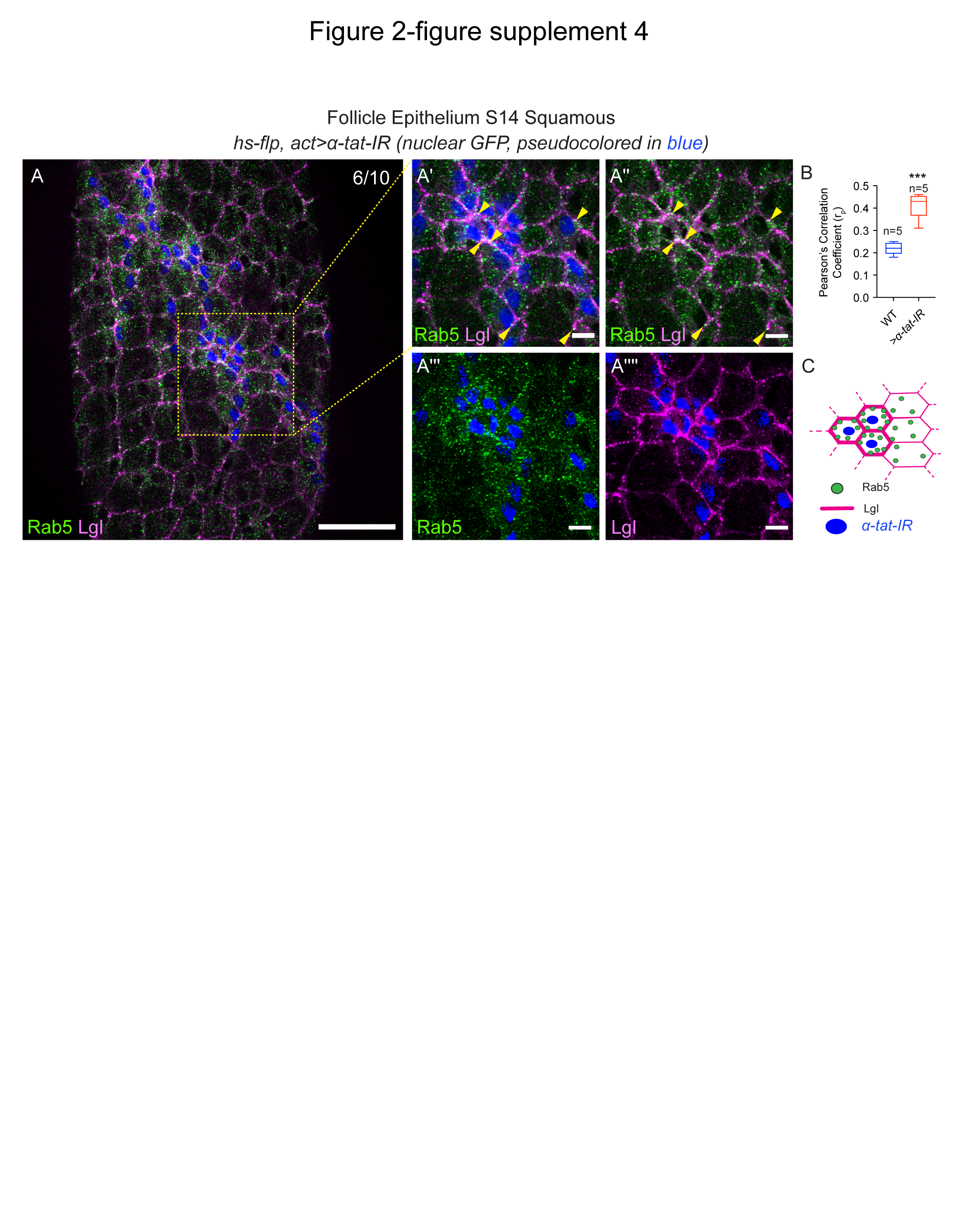

### Figure 3-figure supplement 1

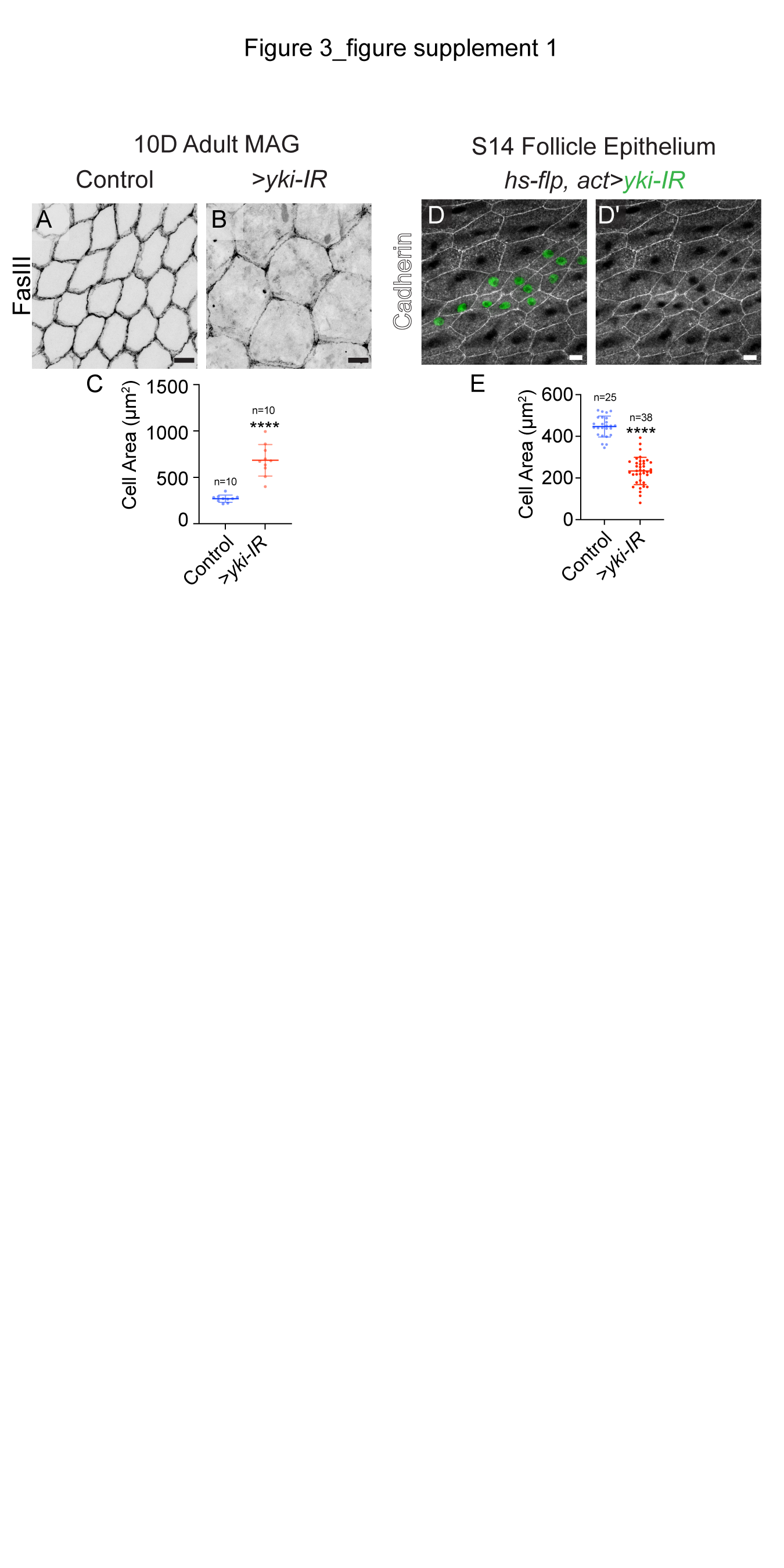

### Figure 3-figure supplement 2

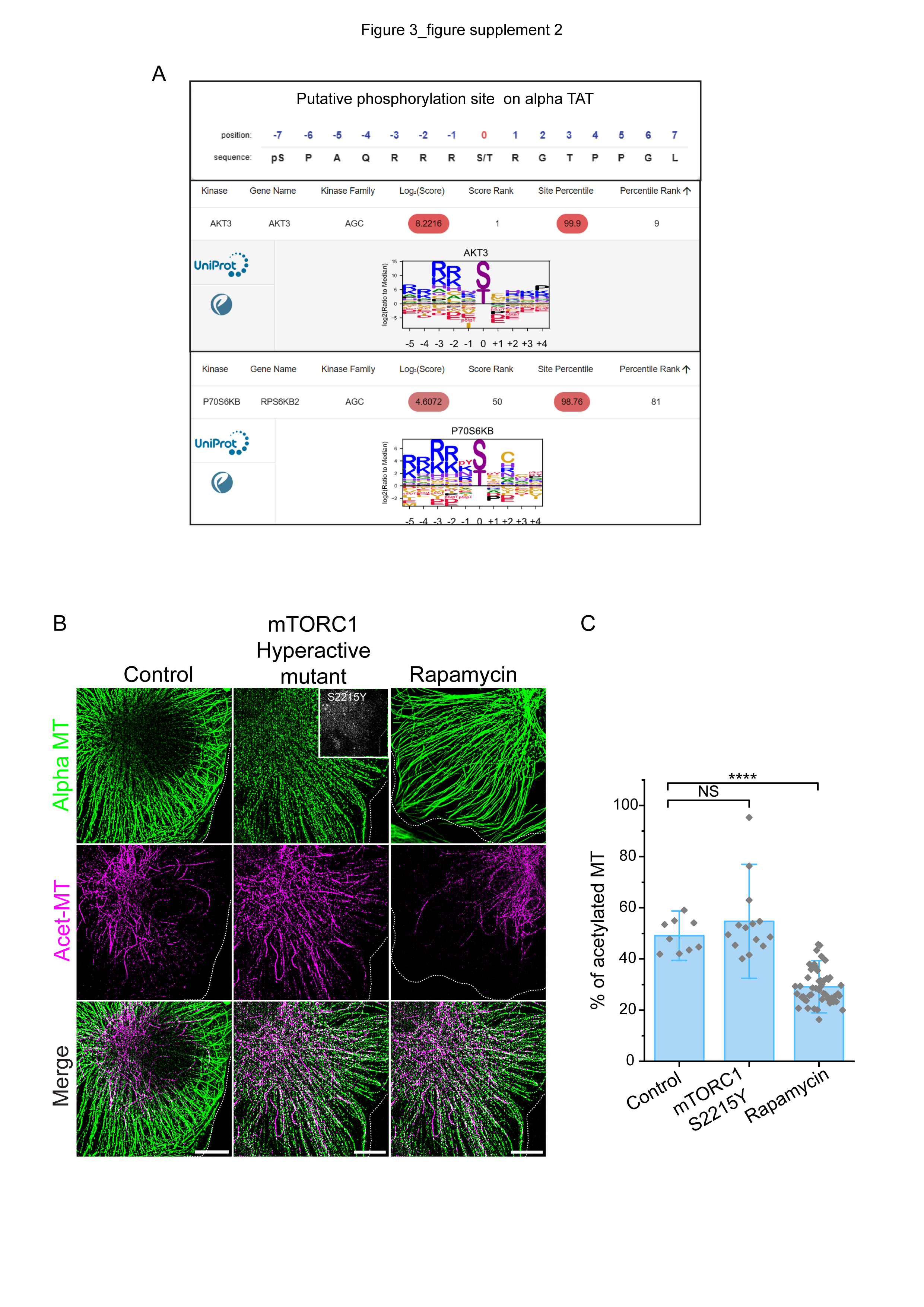
